# Defining biologically directed therapy target signatures in urothelial carcinoma: A transcriptomic framework for precision therapy

**DOI:** 10.64898/2026.06.24.734314

**Authors:** Edwin Lin, Bing-Jian Feng, Kaniz Fatema, Zeynep Irem Ozay, Georges Gebrael, Varun Nandakumar, Ethan Murdock, Haoran Li, G. Daniel Grass, Eric Singer, Laura Graham, Qiang Li, Bodour Salhia, Saum Ghodoussipour, Jennifer King, Ken Nepple, Zin Myint, Paul Viscuse, Michelle Churchman, David Lum, Umang Swami, Neeraj Agarwal, Sumati Gupta

## Abstract

**Introduction:** Nectin-4 targeting antibody-drug conjugate (ADC) enfortumab vedotin (EV), in combination with pembrolizumab, is the first-line treatment for patients with locally advanced or metastatic urothelial carcinoma (UC). Optimal treatment strategies for patients who are non-responders or progress on EV with pembrolizumab remain an unmet clinical need. We sought to characterize ADC and immunotherapy (IO)-associated target expression profiles to identify candidate therapeutic vulnerabilities beyond EV.

**Methods:** We conducted a literature review to identify ADC and IO targets with approved or investigational relevance in UC. Unsupervised hierarchical clustering was used to identify clusters of target gene expression in RNA-seq data. Transcriptomic clustering analyses were performed in 434 patients from The Cancer Genome Atlas Bladder Urothelial Carcinoma cohort (TCGA-BLCA) and validated in an independent cohort of 478 patients from the Oncology Research Information Exchange Network (ORIEN) consortium. Proteomic interrogation of these targets was performed using mass spectrometry data from additional cohort of 116 patients. Differential gene expression analyses evaluated associations between target expression patterns, histologic variants, and consensus molecular subtypes of muscle-invasive bladder cancer (CMIBC).

**Results:** We identified 13 ADC and 10 IO-associated targets with translational relevance in UC. Transcriptomic analyses revealed three reproducible clusters of overexpressed target genes across independent cohorts: 1) a luminal/epithelial-associated cluster enriched for *VTCN1, SLITRK6, FGFR3, NECTIN4, TACSTD2, ERBB2,* and *ERBB3;* 2) an immune target predominant cluster enriched for *BTLA, LAG3, PDCD1, TIGIT, CTLA4, TNFRSF9, TNFRSF18, TNFRSF4*; and 3) a basal/neuroendocrine-associated cluster characterized by *CD274, F3, NT5E, EGFR, MET* and *DLL3.* Similar clusters were largely conserved at the proteomic level. Adenocarcinomas overexpressed *ERBB3* compared to neuroendocrine and squamous cell carcinomas. Pure squamous cell carcinomas overexpressed *TACSTD2* compared to adenocarcinomas. In CMIBC subtypes, basal/squamous tumors expressed higher levels of *CD274, EGFR*, *F3*, *LAG3*, *NT5E*, and *TNFRSF18*, whereas luminal tumors demonstrated higher ERBB2 and ERBB3 expression. Neuroendocrine-like tumors showed higher *DLL3* expression compared to all other subtypes. Tumors with low expression of *NECTIN4*, *TACSTD*2, and *FGFR3* were enriched for alternative targets including *DLL3, CD274,* and *CD276.* Our findings provide a framework for hypothesis-driven therapeutic prioritization in advanced UC.

Conclusions:

UC is characterized by reproducible, biologically distinct patterns of ADC and IO target expressions. The degree of expression of *NECTIN4* was positively associated with *TACSTD2*, *FGFR3* and inversely associated with *DLL3*, *CD276*, and *CD274*, supporting alternative biologically informed treatment strategies besides EV . Histologic variants and molecular subtypes of UC also display distinct patterns of target expression. This study provides the first integrated transcriptomic framework linking ADC and IO target co-expression patterns for hypothesis-driven therapeutic prioritization. These findings provide a basis for rational ADC and immunotherapy development in advanced UC and support prospective proteomic validation in treatment stratified cohorts.

**Statement of Translational Relevance:** Enfortumab vedotin plus pembrolizumab has redefined first-line therapy for advanced urothelial carcinoma, yet treatment selection following resistance or progression remains undefined. In this study, we integrate transcriptomic and proteomic analyses across independent cohorts to define reproducible patterns of antibody–drug conjugate (ADC) and immunotherapy target co-expression in urothelial carcinoma. We identify biologically distinct target-expression patterns that are associated with histologic and molecular subtypes and demonstrate coordinated and, in some cases, mutually exclusive relationships among therapeutically actionable targets.

These findings have direct translational implications. First, they provide biologic rationale for rational sequencing and combination strategies based on co-expressed targets in *NECTIN4*-enriched tumors. Second, they identify alternative therapeutic vulnerabilities, including *DLL3*- and *CD274*-associated pathways, in tumors with low *NECTIN4* expression, a population potentially enriched for resistance to EV-based therapy. Finally, this framework establishes a foundation for biomarker-driven clinical trials in urothelial carcinoma and supports the development of precision therapeutic approaches beyond current standards.

## Introduction

For decades, first line therapy for locally advanced or metastatic (referred to as advanced hereafter) urothelial carcinoma (UC) comprised platinum-based chemotherapy but associated with limited survival benefit and significant adverse effects. Recent studies have demonstrated superior outcomes with antibody-drug conjugates (ADCs), which selectively deliver cytotoxic payloads to tumor-associated antigens^1^. Together with immune therapies, ADCs have fundamentally transformed the therapeutic landscape across multiple malignancies by improving response rates and in some cases achieving durable clinical benefit compared with conventional chemotherapy^2,3^.

In advanced UC, Nectin-4 targeting ADC, enfortumab vedotin (EV) combined with pembrolizumab has demonstrated superior overall survival (OS), achieving a median OS of 31.5 months compared to 16.1 months with chemotherapy^4^. This combination (henceforth abbreviated as EV plus pembrolizumab) is now the standard first-line treatment for advanced UC ^4^. Despite remarkable outcomes, approximately one-third of patients fail to respond to this treatment. Therefore, there is a vital clinical need for treatment strategies for patients who are non-responders or experience disease progression on EV plus pembrolizumab. ADCs against other molecular targets are under active investigation for UC as well as other solid tumors and may offer viable treatment strategies for these patients. Similarly, immunotherapy targets are also advancing to multiple immune checkpoints and novel bispecific antibodies and may be harnessed for durable responses. However, the expression landscape and co-occurrence patterns of actionable ADC and IO targets in relation to Nectin-4 expression remains poorly characterized (or incompletely defined) in UC.

To address these questions, we utilized transcriptomic profiling to characterize reproducible ADC and IO target expression patterns in UC across independent patient cohorts. Our study defines transcriptionally distinct therapeutic target-expression states in UC and provides a framework for biomarker-guided therapeutic development beyond EV plus pembrolizumab.

## Methods

### Literature Search

Identification of ADC and IO targets relevant to our study was performed using the following literature search methodology: Search terms include “bladder cancer”, “urothelial carcinoma”, “antibody-drug conjugate”, “ADC”, “solid tumor”, and “immunotherapy”. Molecular targets were included for downstream analysis based on whether there were therapeutics targeting these compounds with ongoing clinical trials or had achieved FDA. Targets were not restricted to urothelial carcinoma but included ADC and IO targets with clinical or investigational relevance across solid tumors. Molecular targets for hematologic malignancies were excluded.

### Patient Cohorts

Discovery of ADC and IO target signatures in UC was performed within The Cancer Genome Atlas (TCGA) cohort. TCGA cohort clinical characteristics and methodologies are described in detail in a previous publication ^5^. Briefly, this cohort comprised 406 patients with 412 tumors and 19 matched normal bladder urothelium samples. Validation of ADC and IO patterns in UC was performed using a separate, independent cohort of patients in the Oncology Research Information

Exchange Network (ORIEN) cohort. The ORIEN cohort clinical characteristics are described in a previous publication^6^. The ORIEN cohort comprised tumors from 478 patients, and RNA-seq of matched normal bladder urothelium samples was not performed within this cohort. Proteomic comparison of ADC and IO targets was examined in a previous study by Xu et al ^7^. This dataset comprised a cohort of 116 patients with 157 tumors and paired 76 normal bladder urothelium who underwent proteomic profiling by high-performance liquid chromatography followed by label-free mass spectrometry.

### Bioinformatics

Differential gene expression analysis was performed using DESeq2. Unwanted technical variation was estimated using the RUVSeq package (RUVg, k = 6) with a set of 150 empirically defined stable genes, and the resulting W factors were incorporated as covariates. RUV-normalized counts were variance-stabilized using the DESeq2 (v1.48.2) variance stabilizing transformation (VST) for downstream analyses. RNA-seq differential expression analysis was performed using DESeq2 with a generalized linear model that included CMIBC subtype and covariates (sex, age at specimen collection, stage, sample type, library preparation method (TruSeq vs. Tagmentation), and W factors for unwanted variation). Pairwise contrasts between CMIBC subtypes were computed using Wald tests with adaptive shrinkage of log2 fold changes (ashr), and false discovery rate (FDR) correction was applied across ADC target genes using the Benjamini–Hochberg method. Criteria used to determine significance included false discovery rate-adjusted p value less than 0.01 and Log2 fold change with magnitude greater than 1.

Downstream bioinformatic analyses were performed using Python 3.13.5. Fisher’s exact test was used to compare categorical clinical features between the two clusters. Survival analysis was performed using the Log-Rank test. Hierarchical clustering was performed with scipy 1.15.3 using the Ward method on log-scaled TPM. Pearson correlation coefficients for ADC target gene expression was calculated using scipy 1.15.3. Dendrograms and heatmaps were plotted using seaborn 0.13.2.

### Multiplex Immunohistochemistry

Multiplex immunohistochemistry (mIHC) and immunofluorescence analyses were performed on selected malignant ascites clinical specimens to evaluate target co-expression patterns. Detailed experimental procedures, antibody conditions, and image analysis workflows are described in the Supplementary Methods.

## Results

### Identification of therapeutically relevant ADC and immunotherapy targets

A systematic literature review identified 23 therapeutically relevant targets under active investigation or with FDA approval for solid tumors, including 13 ADC and 10 IO-associated targets (**Table 1)**.

**Table 1.**

| *Target Gene* | *Protein ID* | *Tumor type* | *Investigational status* |
| --- | --- | --- | --- |
| *BTLA* | BTLA | ESCC<sup>17</sup><br>NSCLC/SCLC<sup>18</sup> | IND |
| *CD274* | PD-L1 | UC<sup>4,19-21</sup> | FDA approved |
| *CD276* | B7-H3 | NSCLC/SCLC<sup>22</sup> | Clinical Trial |
| *CEACAM5* | CEACAM5 | NSCLC<sup>23</sup><br>CRC<sup>24</sup><br>Pan-Tumor<sup>25</sup> | Clinical Trial |
| *CTLA4* | CTLA-4 | UC<sup>20</sup> | Clinical Trial |
| *DLL3* | DLL3 | SCLC<sup>26</sup> | Clinical Trial |
| *EGFR* | EGFR | GBM<sup>27</sup><br>NPC<sup>28</sup> | Clinical Trial |
| *ERBB2* | HER2 | Breast<br>Gastric<sup>29</sup><br>NSCLC<sup>30</sup><br>Solid tumors<sup>31</sup><br>UC<sup>32</sup> | FDA approved |
| *ERBB3* | HER3 | Breast<sup>33</sup><br>NSCLC<sup>34</sup> | Clinical Trial |
| *F3* | TF | Cervical<sup>35</sup> | FDA approved |
| *FGFR3* | FGFR3 | UC<sup>36</sup> | Pre-Clinical |
| *LAG3* | LAG3 | Melanoma<sup>37</sup> | Clinical Trial |
| *MET* | MET | NSCLC<sup>38</sup> | FDA approved |
| *NECTIN4* | NECTIN4 | UC<sup>4</sup> | FDA approved |
| *NT5E* | CD73 | Solid Tumors<sup>39</sup> | Clinical Trial |
| *PDCD1* | PD-1 | UC<sup>4,19-21</sup> |  |
| *SLITRK6* | SLITRK6 | UC<sup>40</sup> | Clinical Trial |
| *TACSTD2* | TROP2 | UC<sup>41</sup> | Clinical Trial |
| *TIGIT* | TIGIT | NSCLC, SCLC, Cervical<br>cancer<br>Pan-Tumor<sup>42</sup> | Clinical Trial |
| *TNFRSF18* | GITR | Pan-Tumor | Clinical Trial |
| *TNFRSF4* | OX40L | Solid Tumors<sup>43</sup> | Clinical Trial |
| *TNFRSF9* | 4-1BB | Solid Tumors<sup>44</sup> | Clinical Trial |
| *VTCN1* | B7-H4 | Solid Tumor<sup>45</sup> | Clinical Trial |

### Reproducible transcriptomic target-expression states across independent cohorts

Unsupervised Hierarchical clustering of transcriptomic data from the TCGA-BLCA discovery cohort identified three major clustering patterns of target co-expression (**Figure 1A**). Cluster 1 included *VTCN1, SLITRK6, FGFR3, NECTIN4, TACSTD2, ERBB2,* and *ERBB3.* Cluster 2 included *BTLA, LAG3, PDCD1, TIGIT, CTLA4, TNFRSF9, TNFRSF18, TNFRSF4*. Cluster 3 included *CD274, F3, NT5E, EGFR, MET, DLL3*.

**Figure 1.**
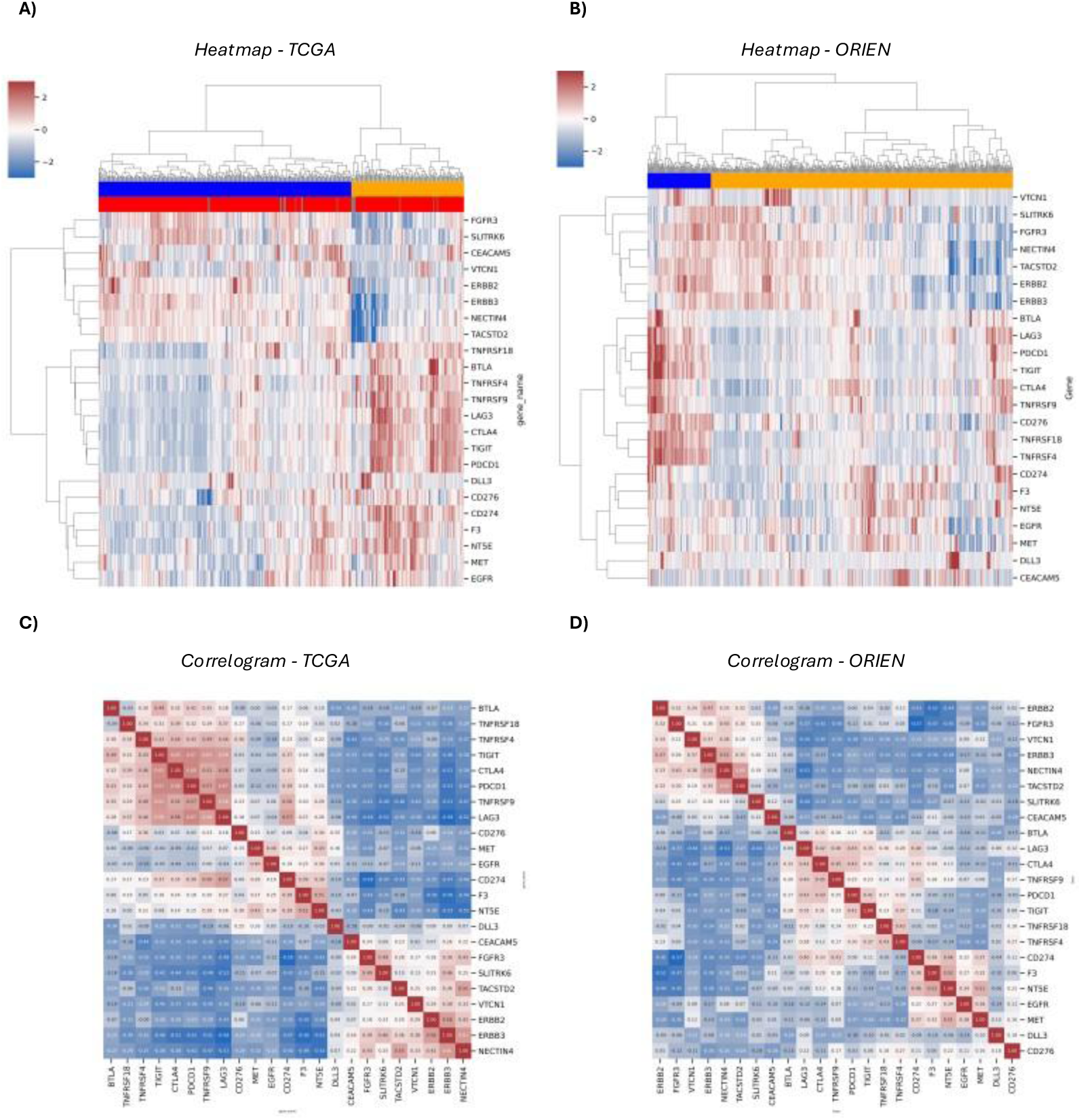
Transcriptomic profiles of target expression in UC. A) Discovery cohort (TCGA). B) Validation cohort (ORIEN). Heatmaps were plotted with log-transformed Z-score of transcripts per million for each gene. Row dendrograms represent gene clusters and column dendrograms represent tumor subsets. Pearson s correlation between ADC and IO targets in UC are plotted in a heatmap based on log-transformed Z-score of transcripts per million for C) TCGA and D) ORIEN cohorts.

These findings were validated in the ORIEN cohort, which demonstrated highly concordant clustering patterns (**Figure 1B**). Minor differences in cluster composition were observed (including *CD276* and *CEACAM5*), but overall architecture was preserved. Minor differences in cluster composition were observed (including *CD276* and *CEACAM5*), but overall architecture was preserved.

Given the central role of Necctin-4 as a therapeutic target, we examined the relationship of *NECTIN4* expression with other ADC and immunotherapy-associated genes (**Figure 1C, 1D**).. *NECTIN4* expression demonstrated positive correlations with luminal ADC-associated targets, including *FGFR3*, *ERBB2*, and *TACSTD2* and inverse associations with basal/neuroendocrine and immune-associated targets, including *EGFR, MET,* and *CD274* These findings define biologically distinct and potentially divergent target-expression states in UC.

### Partial proteomic concordance supports biological relevance of target-expression states

To evaluate concordance of the clusters at a protein level, we analyzed mass spectrometry data within a previous study comprising 116 patients who underwent label-free mass spectrometry profiling (**Supplementary Figure 1**). A total of 13 out of 23 targets were quantifiable proteomically, and hierarchical clustering of this subset yielded similar clusters consistent with transcriptomic analyses.

### Limited association of target-expression states with clinical outcomes

Tumor-expression clusters were not significantly associated with any disease stage (**Table 2**, Fisher’s Exact p=0.145), or overall survival (**Supplementary Figure 2**). Similarly, expression of individual ADC and IO targets did not show significant associations with overall survival that could be replicable across both cohorts (Supplementary **Figure 3, 4**). These analyses are therefore considered exploratory and not used for downstream inference.

**Table 2.**

| *Stage* | *Cluster 1* | *Cluster 2* |
| --- | --- | --- |
| *Stage IV* | 86 | 35 |
| *Stage III* | 78 | 50 |
| *Stage II* | 77 | 29 |
| *N/A* | 53 | 19 |
| *Stage I* | 4 | 0 |

### Histologic subtypes exhibit distinct target-expression patterns

Histologic variants of urothelial cancer may have differing alterations and treatment response to treatment ^10^. Hence, we evaluated histology-specific patterns in ADC and IO target expression in the ORIEN cohort, which revealed differential expression of ADC and immunotherapy targets across histologic variants (Figure 2, Supplementary Table 1). Key observations included:

- Conventional urothelial carcinomas demonstrated higher expression of *NECTIN4, FGFR3, ERBB2,* and *TACSTD2* relative to neuroendocrine and adenocarcinoma subtypes
- Squamous cell carcinomas were enriched for *EGFR, F3, NECTIN4,* and *TACSTD2*
- Adenocarcinomas demonstrated increased *ERBB3* expression
- Neuroendocrine tumors showed increased *DLL3* expression, although this did not reach statistical significance, likely due to limited sample size

**Figure 2.**
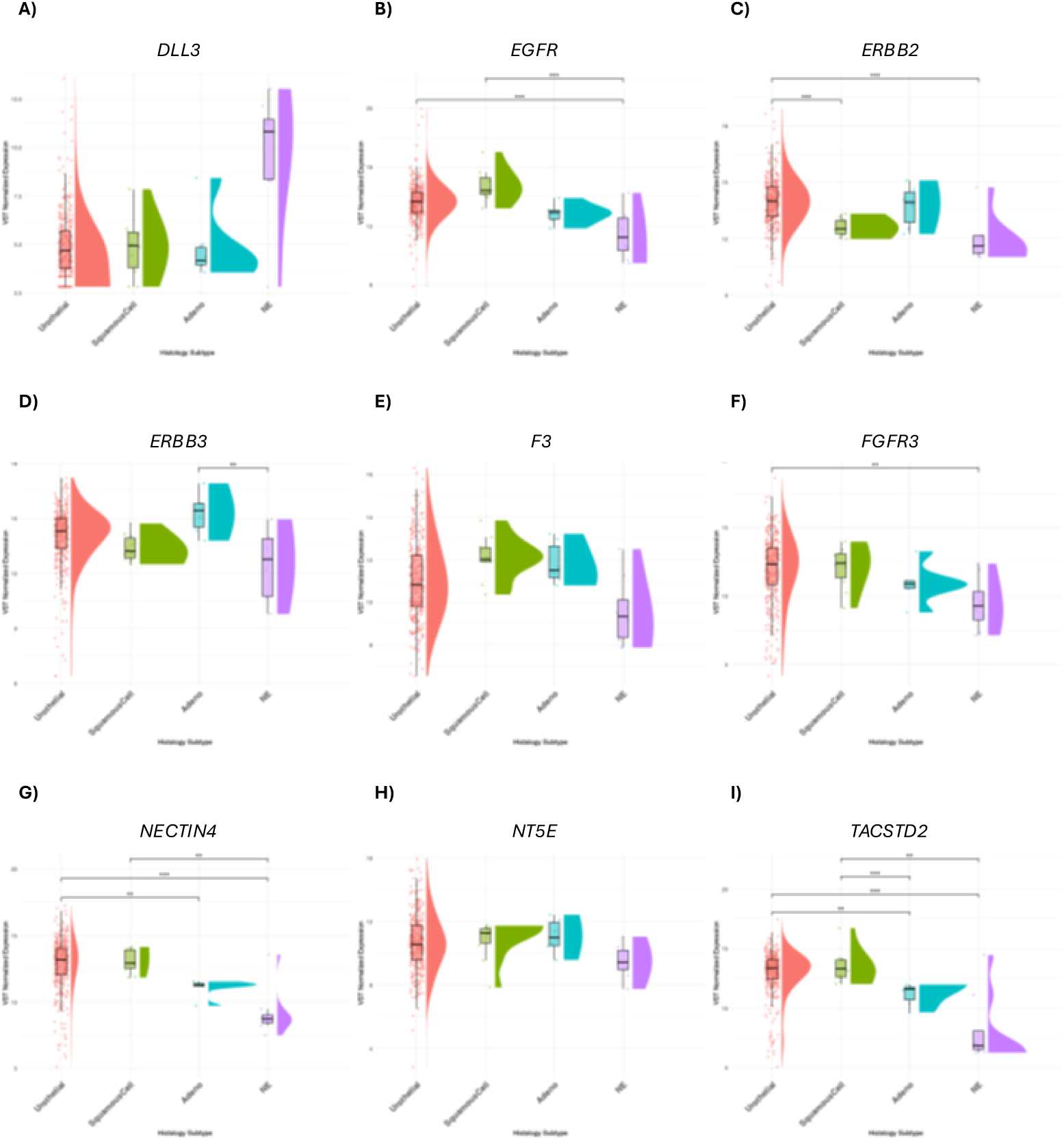
Differentially expressed ADC and IO target genes by bladder carcinoma histologic type. Variance stabilized counts were plotted according to histologic subtype. Genes that were significantly differentially expressed between at least one comparison were included. Brackets denote significant differential expression between two cohorts, as defined by Log2(Fold Change) 1 and false discovery rate adjusted P-value 0.01.

These findings highlight heterogeneity in target expression across histologic subtypes.

UC with squamous or glandular differentiation are often inconsistently described in pathology reports^11^, but known to be significantly more common than pure adenocarcinoma or squamous cell carcinoma of the bladder. Our manual curation of pathology reports in both TCGA and ORIEN cohorts was not only reflective of this but was also notable for UC with varying degrees of small cell differentiation. We were unable to achieve granularity in the relative composition of histologic variants in these retrospective datasets.

### Target-expression states align with molecular subtypes of urothelial carcinoma

We next evaluated associations between ADC and IO target expression molecular subtypes of urothelial carcinoma according to CMIBC transcriptomic classification^12^ (Figure 3, Supplementary Table 2).

- Basal/squamous tumors demonstrated increased expression of *CD274, EGFR, F3, LAG3,* and *NT5E*
- Luminal tumors were enriched for *ERBB2* and *ERBB3*
- Neuroendocrine-like tumors exhibited increased *DLL3* expression

**Figure 3.**
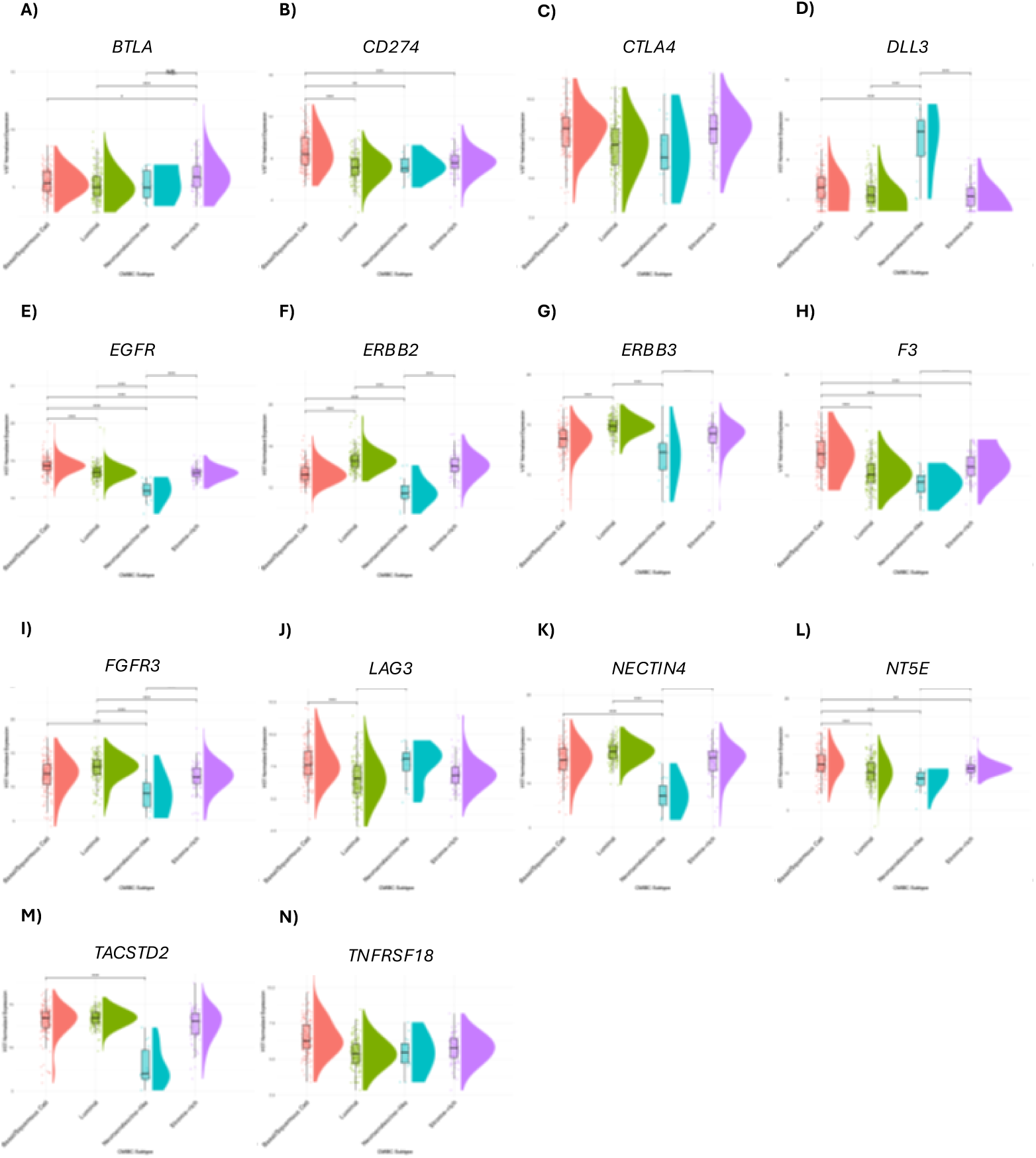
Differentially expressed ADC and IO target genes by molecular subtype of bladder carcinoma. Variance stabilized counts were plotted according to CMIBC subtype. Genes that were significantly differentially expressed between at least one comparison were included. Brackets denote significant differential expression between two cohorts, as defined by Log2(Fold Change) 1 and false discovery rate adjusted P-value 0.01.

These findings demonstrate concordance between transcriptomic target-expression states and established molecular classification frameworks.

### Clinical exemplars illustrate biologically informed target prioritization

To illustrate the potential translational relevance of these findings, we present representative clinical cases.

A patient with metastatic neuroendocrine carcinoma demonstrated a durable response to DLL3-targeted therapy following progression on platinum-based chemotherapy and immunotherapy (Figure 4A–B). This observation is consistent with enrichment of DLL3 expression within the neuroendocrine-associated target-expression state identified in cohort analyses.

**Figure 4.**
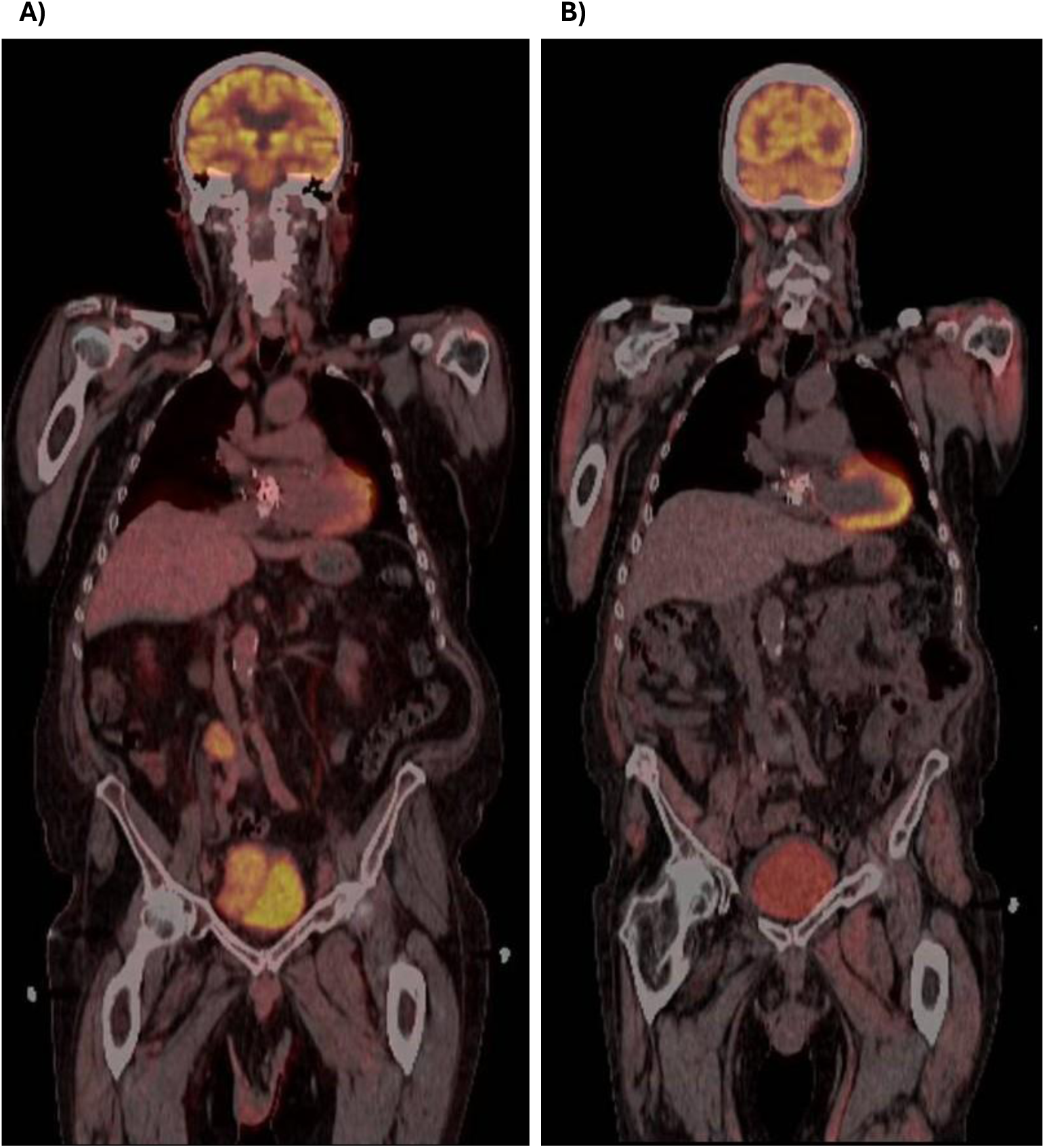
PET-CT for a UC patient A) At the time of disease progression after partial response to carboplatin/etoposide with atezolizumab. B) PET-CT after treatment with tarlatamab-dlle

In a second case, a patient with metastatic urothelial carcinoma progressing after multiple lines of therapy, including enfortumab vedotin and sacituzumab govitecan, underwent multiplex immunohistochemistry of malignant ascites. This demonstrated persistent co-expression of NECTIN4, FGFR3, HER2, and TROP2, despite prior exposure to ADCs targeting these antigens. These findings mirror the luminal ADC-associated co-expression patterns observed in transcriptomic analyses (Figure 5B–C; Supplementary Figure 5).

**Figure 5.**
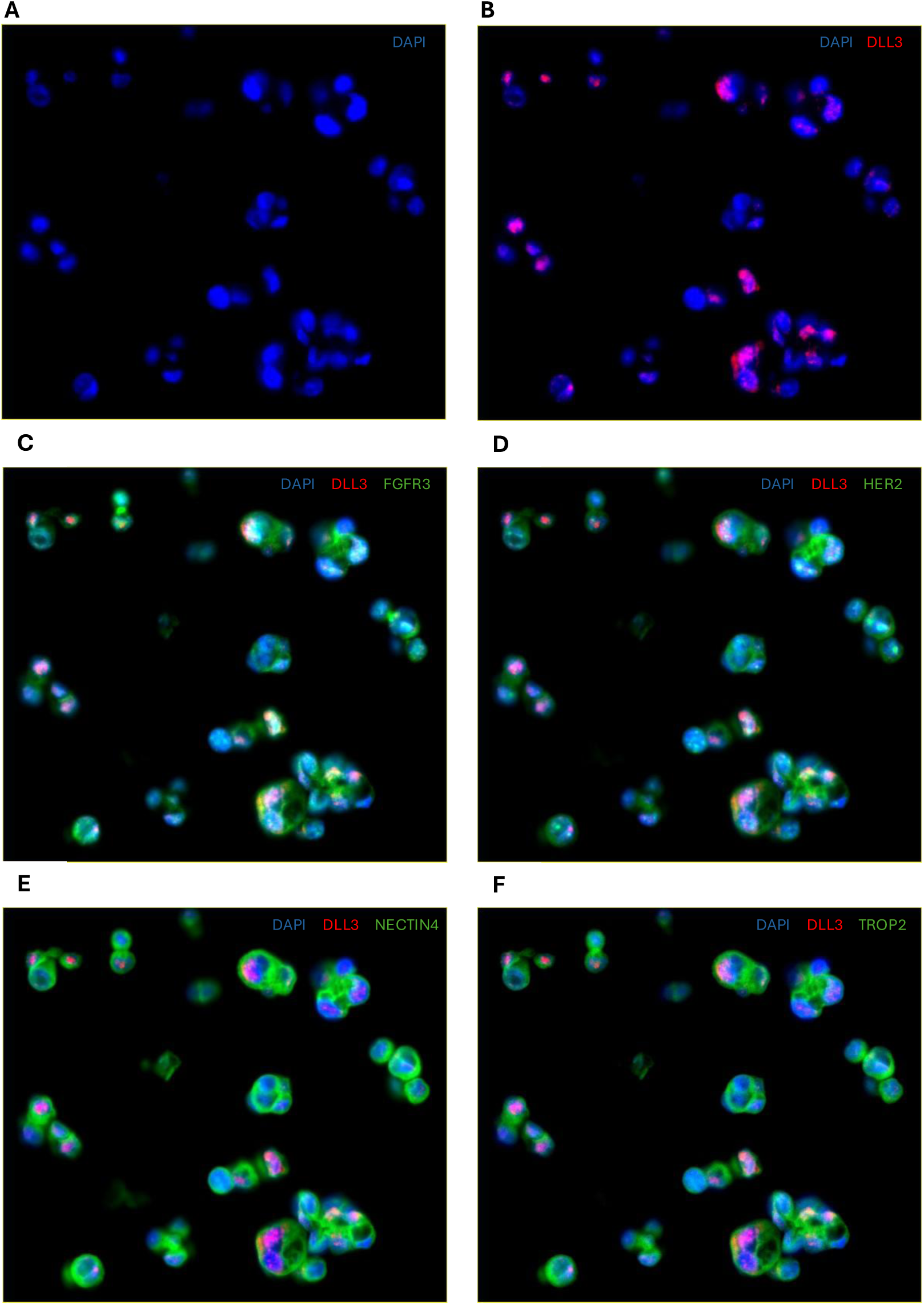
Multiplex immunohistochemistry of malignant ascites for a UC patient after treatment progression with enfortumab vedotin and sacituzumab govetecan. DAPI is represented in blue. DLL3 is represented in red. Given the similar pattern of cytoplasmic and membranous staining with A) FGFR3, B) HER2, C) Nectin-4, and D) Trop-2 these are represented in green and shown in separate figures.

These cases are illustrative and hypothesis-generating, supporting the biological plausibility of target-expression–guided therapeutic strategies but not establishing clinical efficacy. These findings establish a conceptual framework for biologically informed therapeutic prioritization in UC, particularly in the setting of resistance to EV-based therapy. These observations are hypothesis-generating and require prospective validation.

### Integrated framework defines biologically distinct therapeutic target-expression states

The schematic in Figure 6 illustrates a biologically informed therapeutic framework derived from integrated transcriptomic and proteomic analyses across independent urothelial carcinoma (UC) cohorts. Three reproducible patterns of target co-expression were identified: (1) a luminal/epithelial-associated ADC cluster enriched for *NECTIN4, FGFR3, ERBB2, ERBB3,* and *TACSTD2*; (2) an immune checkpoint–enriched cluster characterized by expression of *PDCD1, CTLA4, LAG3,* and *TIGIT*; and (3) a basal/neuroendocrine-associated cluster enriched for *EGFR, MET, CD274,* and *DLL3*.

**Figure 6.**
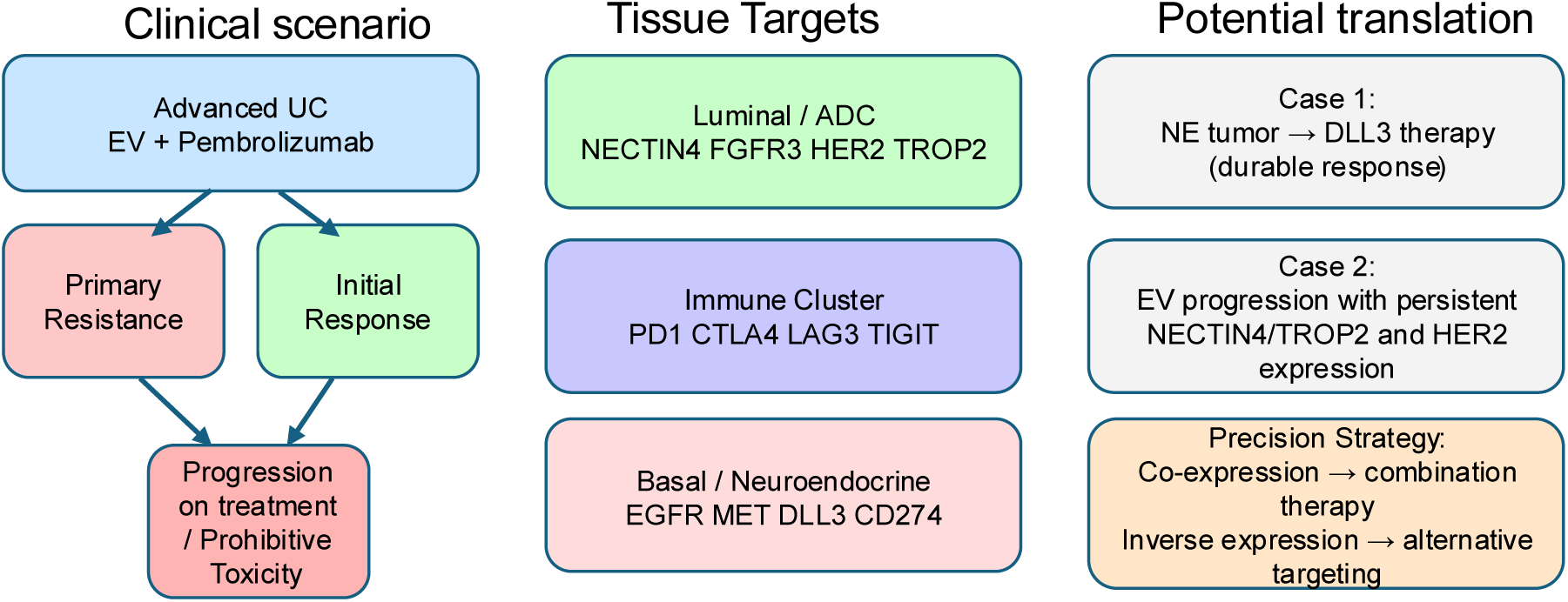
Transcriptomic framework for biologically guided selection of antibody -drug conjugate (ADC) and immunotherapy targets in urothelial carcinoma.

The framework highlights both co-occurring and mutually exclusive target expression relationships, with *NECTIN4* expression positively correlated with luminal ADC-associated targets and inversely associated with several immune and basal/neuroendocrine targets, including *CD274, EGFR, MET,* and *DLL3*.

These patterns define clinically relevant therapeutic strategies across distinct biological contexts:

- *NECTIN4*-high tumors may benefit from combination ADC strategies targeting co-expressed luminal markers (e.g., *FGFR3*, *HER2*, *TROP-2*).
- *NECTIN4*-low tumors demonstrate enrichment of alternative targets, including *DLL3*, *MET*, *EGFR*, *CD274*, and *CD276*, supporting therapeutic prioritization beyond enfortumab vedotin.
- Neuroendocrine-like tumors exhibit preferential *DLL3* expression, providing a rationale for DLL3-directed therapies.
- Immune-enriched tumors demonstrate upregulation of multiple immune checkpoints beyond PD-1/PD-L1, suggesting potential responsiveness to combinatorial immunotherapy approaches.

Together, this framework provides a conceptual model for precision therapeutic selection based on tumor-specific target expression states, particularly in patients with primary resistance or progression following enfortumab vedotin plus pembrolizumab. These findings are hypothesis-generating and support prospective validation in treatment-annotated cohorts.

## Discussion

In this study, we define a reproducible and biologically coherent landscape of ADC and immunotherapy target expression in urothelial carcinoma, identifying three principal patterns of co-expression that are conserved across independent transcriptomic cohorts and supported by orthogonal proteomic data. These findings move beyond single-target analyses to establish coordinated target-expression states, providing a conceptual framework for therapeutic prioritization in UC.

The clinical relevance of this work is underscored by the evolving treatment paradigm in advanced UC. Although EV plus pembrolizumab has significantly improved outcomes, a substantial proportion of patients experience primary resistance or disease progression. In this setting, treatment selection is not currently guided by tumor biology. Our findings address this gap by defining distinct target-expression states that may inform rational therapeutic strategies following EV-based therapy.

A key observation from our analysis is that *NECTIN4* expression, central to EV activity, is not uniformly distributed across tumors and is embedded within a broader network of co-expressed ADC-associated targets, including *FGFR3*, *ERBB2*, and *TACSTD2*. These findings provide biologic rationale for combinatorial or sequential ADC approaches targeting co-expressed antigens in Nectin-4-high tumors. EV plus pembrolizumab has become the first line therapy for patients with advanced UC and was recently incorporated into treatment paradigms for patients with cisplatin-ineligible, muscle invasive UC^13^. Given the possibility of achieving a complete response by adding therapies based on co expression patterns may inform future strategies aimed at improving organ preservation. Early-phase clinical studies combining FGFR3- and Nectin-4-directed therapies have demonstrated promising activity with a remarkable 93.3% overall response rate^14^, suggesting synergy with regards to targeting multiple positively-associated targets in UC. the potential relevance of these co-expression patterns.

Conversely, we identify inverse relationships between the expression of *NECTIN4* and a subset of non-luminal targets EGFR, MET, and DLL3. This suggests that tumors with low *NECTIN4* expression may preferentially rely on alternative biologic programs, potentially explaining mechanisms of resistance to EV-based therapy. These observations generate the hypothesis that therapeutic strategies targeting inversely associated pathways may represent rational approaches in EV-refractory disease.

Our study also identifies inverse associations between *NECTIN4* expression with IO targets including *CD274* (PD-L1). Previous meta-analyses have demonstrated ORR of 68% with EV plus pembrolizumab compared to ORR of 43% with EV monotherapy for metastatic urothelial carcinoma^15^. Interestingly, the proportions of tumors with higher expression of *NECTIN4* and immune checkpoint targets in our cohorts were comparable.

Beyond global clustering, we demonstrate that these target-expression patterns are closely linked to histologic and molecular subtypes of UC. More specifically pertaining to histopathology, the current reporting standard for urothelial carcinoma based on the latest College of American Pathologist synoptic categorizes urothelial carcinoma based on whether portions of the tumor demonstrate other histologic subtypes including squamous cell, adenocarcinoma, neuroendocrine, and sarcomatoid differentiation^16^. Basal/squamous and neuroendocrine-like tumors exhibit enrichment of distinct targets, including *EGFR*, *CD274*, and *DLL3*, whereas luminal tumors are characterized by higher expression of *ERBB2* and *ERBB3*. These findings are particularly relevant given the known heterogeneity of UC and the limitations of current pathology reporting, which often incompletely captures variant histology. Molecularly defined target states may therefore provide a complementary approach to histologic classification for therapeutic decision-making.

The translational relevance of this framework is illustrated through representative clinical cases demonstrating biologically guided target selection. While anecdotal, these examples serve as proof-of-concept that alignment between tumor biology and therapeutic targeting may yield clinically meaningful responses, particularly in rare or treatment-refractory disease contexts.

Several limitations of this study warrant consideration. First, our analyses are primarily based on transcriptomic data, with limited but supportive proteomic validation. Second, both TCGA and ORIEN cohorts predate the widespread use of EV plus pembrolizumab, precluding direct assessment of predictive biomarker utility in the contemporary treatment landscape. Third, survival analyses did not yield consistent associations across datasets, likely reflecting heterogeneity in cohort composition and treatment exposure. Finally, the clinical examples presented are illustrative and do not establish efficacy.

Accordingly, our findings require prospective validation in treatment-annotated cohorts. Future studies incorporating longitudinal sampling and integration of genomic, transcriptomic, and proteomic data will be essential to define predictive biomarkers and optimize sequencing strategies.

In conclusion, we define a reproducible transcriptomic framework of ADC and immunotherapy target co-expression in UC, identifying biologically distinct target-expression states that are associated with histologic and molecular subtypes. These data provide a foundation for biomarker-driven clinical investigation and rational therapeutic development beyond EV plus pembrolizumab, particularly in the setting of treatment resistance

## Declarations

Ethics approval and consent to participate: The research presented in this study was approved by the University of Utah IRB.

Consent for publication: All authors have consented to publication

Availability of data and material: The data from TCGA used in this study are accessible under project ID TCGA-BLCA and are available at: https://gdc.cancer.gov. The data from the ORIEN consortium that support the findings of this study are not openly available due to reasons of sensitivity and are available from the corresponding author upon reasonable request.

## Authors’ contributions

Conceptualization: EL SG, NA, US

Methodology: EL, BJF, SG, KF Software: EL, BJF

Validation: EL, BJF Formal analysis: EL, BJF Data Curation: EL, BJF, KF

Review & Editing: All authors contributed Visualization: EL, BJF

## Acknowledgements

We acknowledge support from the Biorepository and Molecular Pathology Shared Resource and HCI’s National Cancer Institute Cancer Center Support Grant P30CA042014

## Supplementary Methods

### Multiplex Immunohistochemistry

CD45 -positive cells were depleted from patient ascites samples with EasySep™ Human CD45 Depletion Kit II (STEMCELL Technologies, # 17898) and EasySep™ Magnet (STEMCELL Technologies, # 18000) according to the manufacture protocol. CD45 depleted ascites cells were fixed with 10% Neutral Buffered Formalin (NBF) at room temperature for 2 hours. The fixed samples were collected by low-speed centrifugation at 100 x g for 1 minute, and the NBF solution was discarded. The fixed sample pellet was resuspended in 200uL of Epredia HistoGel™ Specimen Processing Gel (Fisher Scientific, #22-110-678) and allow to solidify into disc shape.

The histogel disc was then placed into a cassette for formalin fixation and paraffin embedding. Formalin-fixed, paraffin-embedded (FFPE) samples were then cut into 5-μm tissue sections.

The sections were processed using Leica BOND automated stainer. Prior to experimental runs, antibody optimization was conducted on control tissues to determine optimal antigen retrieval conditions, primary antibody concentrations, incubation times, and detection chemistries, aiming to preserve archival material and avoid waste. Optimization protocols included single-plex trials, titration series, and signal-to-background evaluations utilizing matched negative controls (isotype and primary-antibody omission).For both immunohistochemistry (IHC) and immunofluorescence (IF) workflows, slides underwent automated on-board deparaffinization, endogenous peroxidase blocking, and heat-induced antigen retrieval using Bond Epitope

Retrieval solutions (ER1, citrate-based, pH ∼6; or ER2, EDTA-based, pH ∼9) based on antibody requirements. For IHC, primary antibodies were incubated on the instrument, followed by polymer-HRP detection, 3,3’-Diaminobenzidine (DAB) chromogen development, and hematoxylin counterstaining. For multiplex IF, primary antibodies were also incubated on the instrument, with the difference that all antibodies and polymer-HRP detection reagents were applied sequentially, followed by Opal fluorophore development (Akoya Biosciences) and DAPI counterstaining as the last step. Appropriate positive and matched negative tissue controls were included in every run to ensure assay reliability. Detailed antibody specifications, including incubation parameters, retrieval conditions, and assigned Opal dyes, are summarized in the **Table (below)**.

| Antibody | Vendor | Product# | Clone | Retrival | Dilution | Incubation time | Opal |
| --- | --- | --- | --- | --- | --- | --- | --- |
| DLL3 | Abcam | AB229902 | EPR22592-18 | ER2 (pH 9.0), 30 min | 1:100 | 15 min | 480 |
| **FGFR3** | Abcam | AB10651 | Polyclonal | ER2 (pH 9.0),<br>30 min | 1:100 | 20 min | 520 |
| **TROP2** | Abcam | AB214488 | EPR20043 | ER2 (pH 9.0),<br>30 min | 1:6000 | 15 min | 540 |
| **Nectin-4** | Abcam | AB192033 | EPR15613-68 | ER1 (pH 6.0),<br>30 min | 1:600 | 30 min | 570 |
| **HER2** | Abcam | AB134182 | EP1045Y | ER2 (pH 9.0),<br>30 min | 1:3000 | 15 min | 620 |
| **Cytokeratin** | Agilent<br>Dako | M3515 | AE1/AE3 | ER1 (pH 6.0),<br>30 min | 1:500 | 60 min | 650 |

Multiplexed immunofluorescence (mIF) images were analyzed in QuPath (version 0.6.0)^8^. Nucleus segmentation was performed by initially selecting 10 regions of interest for manual annotation followed by nucleus segmentation using the STARDIST algorithm ^9^ at ×20 magnification. STARDIST parameters for this analysis are available at (https://github.com/MarkZaidi/Universal-StarDist-for-QuPath/blob/main/Multimodal%20StarDist%20Segmentation.groovy). Cell borders were obtained by expanding the nuclear outlines by 5 pixels. Single cell data including staining intensity for each channel were exported for further analysis. Thresholds were visually determined by an experienced expert for each marker and further applied to each image for cellular positivity computation

---

**Supplementary Figure 1.**
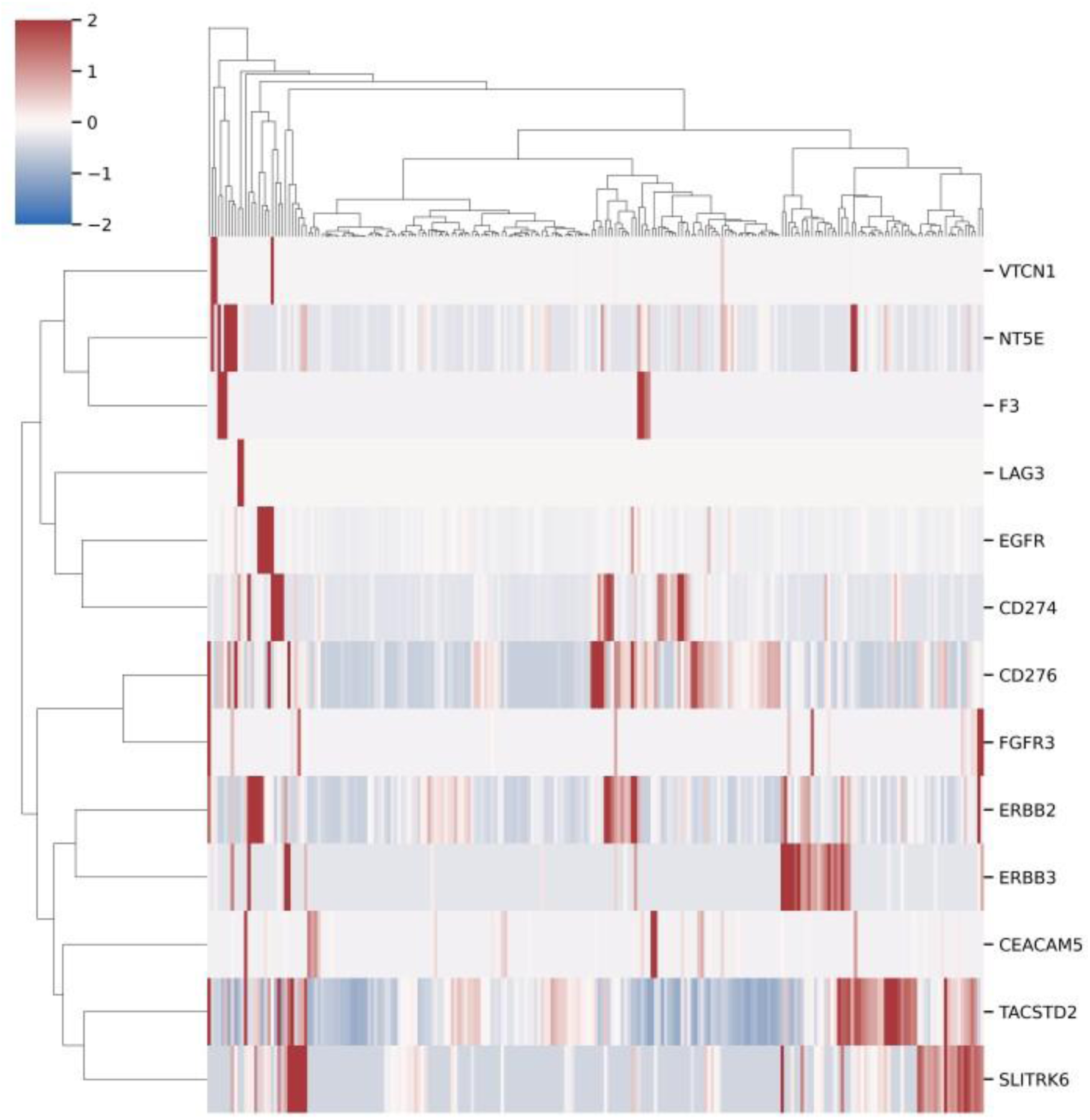
Proteomic clustering of ADC and IO targets in UC. Hierarchical clustering of label-free mass spectrometry data from Xu et al. Rows dendrograms identify clusters of protein expression and column dendrograms represent tumor subsets according to protein expression.

**Supplementary Figure 2.**
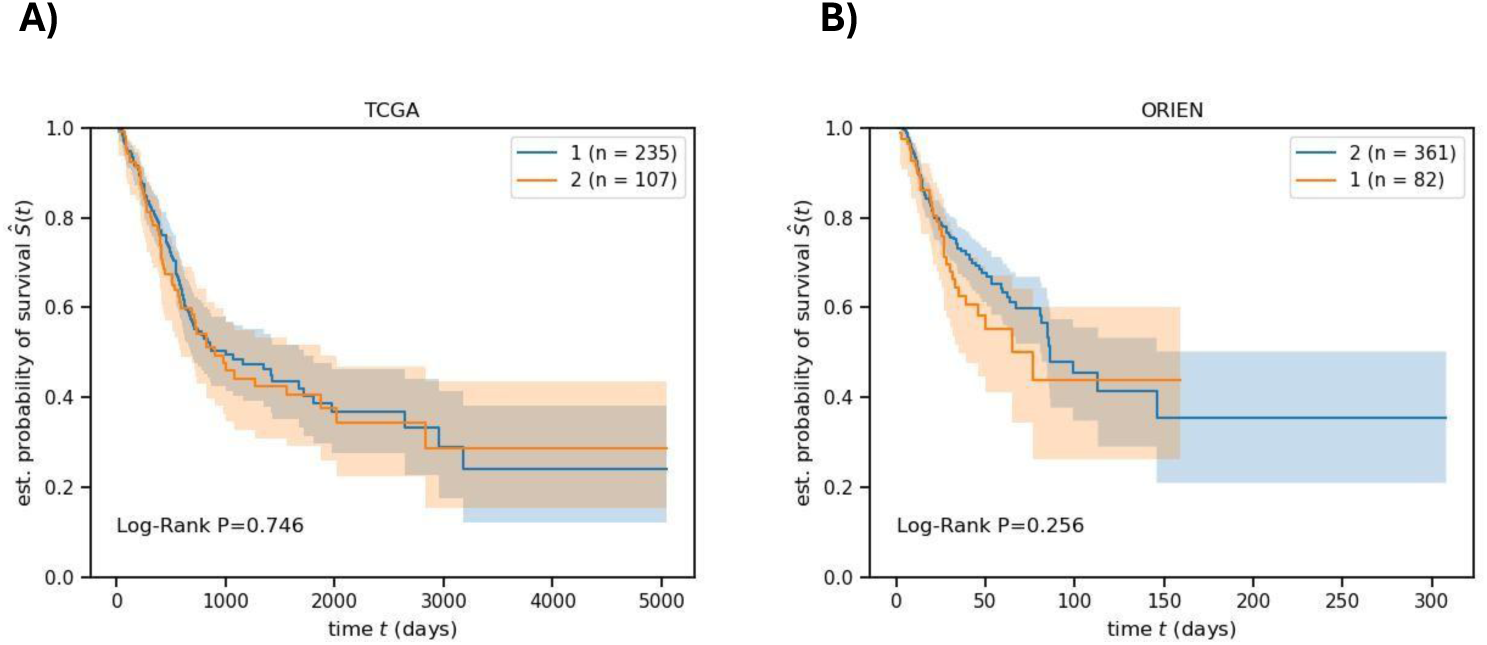
Kaplan-Meier curves of UC in tumor subsets defined by ADC and IO target expression profiles in A) TCGA and B) ORIEN cohorts.

**Supplementary Figure 3.**
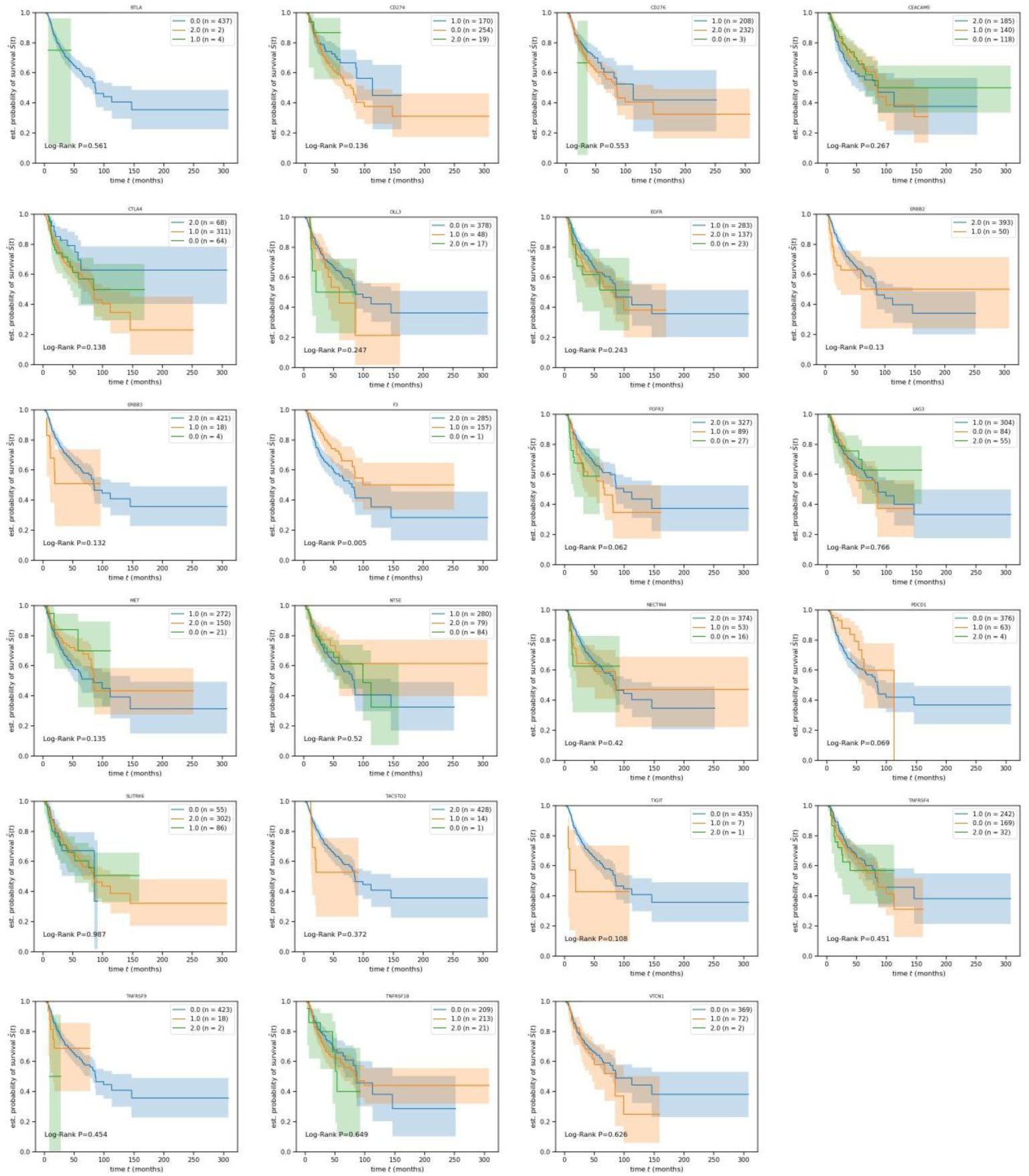
Kaplan-Meier curves of UC according to individual target expression in the ORIEN cohort.

**Supplementary Figure 4.**
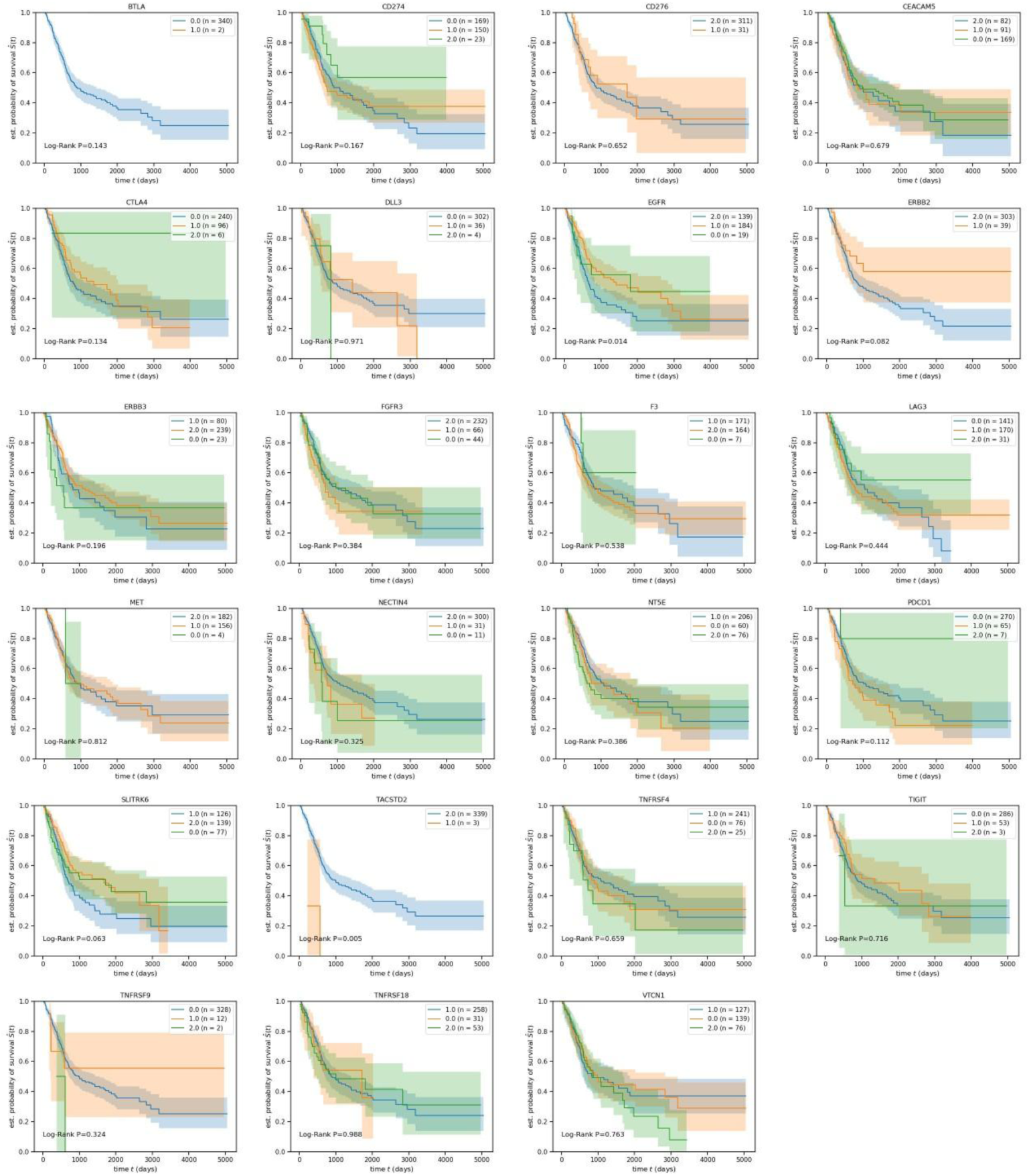
Kaplan-Meier curves of UC according to individual target expression in TCGA cohort.

**Supplementary Figure 5.**
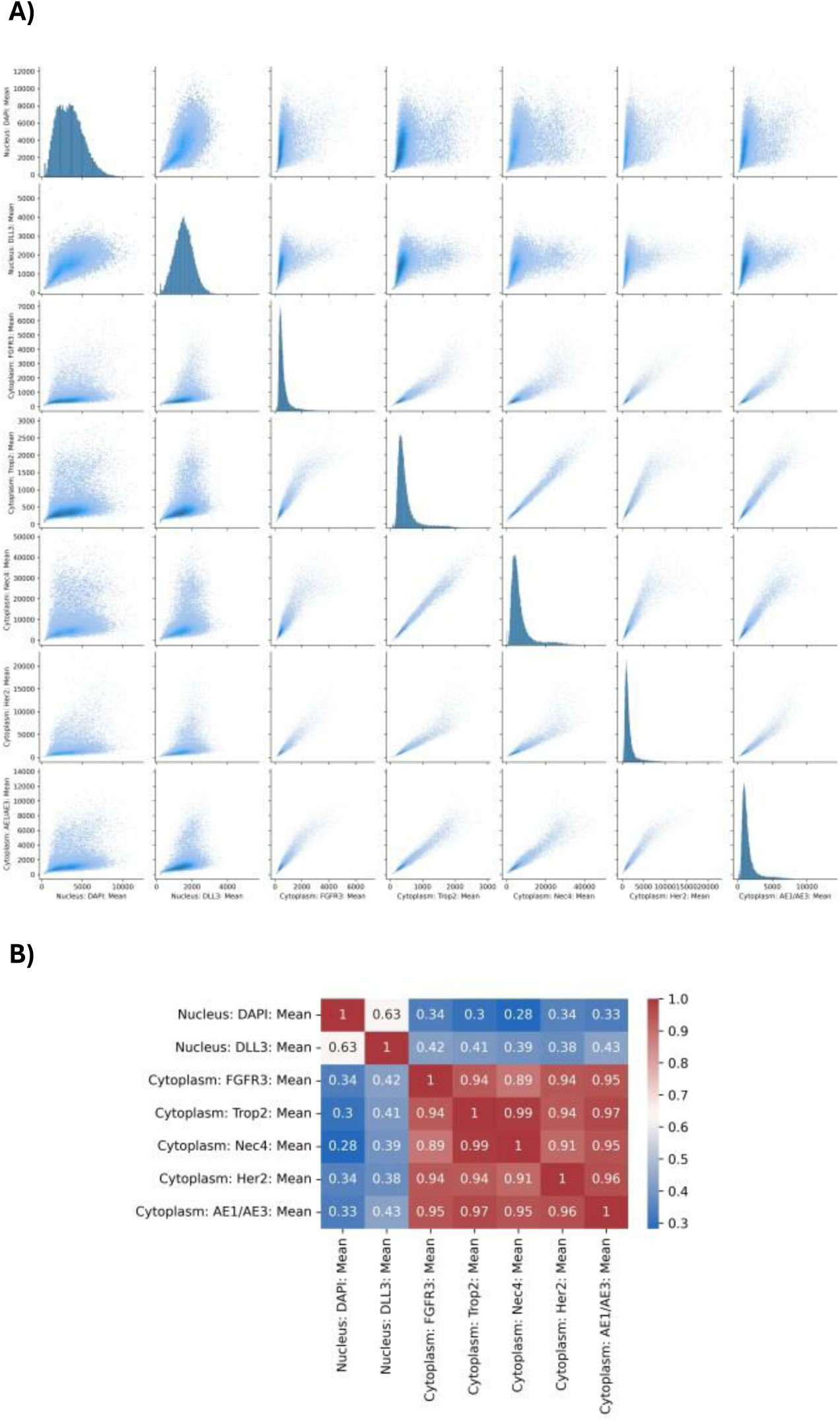
ADC target protein by multiplex immunohistochemistry in a UC patient after progression on enfortumab vedotin and trastuzumab deruxtecan. A) Each scatterplot represents pairwise protein expression denoted on the X and Y axis. Points in each scatterplot represents quantitative intensity of a pair of targets single cells. B) Pearson correlation for each pair of proteins.

